# Targeting C5aR1 Reveals Protective Macrophage Maturation States in Intestinal Injury

**DOI:** 10.1101/2025.11.11.687632

**Authors:** Heather Clark, Ian J. Chai, Stavros Melemenidis, Dhanya K. Nambiar, Kim K. Ryan, Yanyan Jiang, Kerriann M. Casey, Iryna Zherka, Dominika Majorova, Trent M. Woodruff, Edward G. Graves, Quynh-Thu Le, Simon JA. Buczacki, Amato J. Giaccia, Monica M. Olcina

## Abstract

Inflammatory intestinal injury, which may occur following cytotoxic cancer treatments such as radiotherapy, compromises homeostasis when intestinal stem cells undergo apoptosis and regeneration programmes are not effectively engaged. Recent evidence indicates the importance of macrophages in regulating stem regeneration following injury, but how to therapeutically promote specific macrophage subtypes which may tip the balance away from apoptosis and towards regeneration is still unclear. Here we show that targeting complement C5a receptor 1 (C5aR1) reduces radiation-induced intestinal injury by promoting macrophage phenotypes which reduce intestinal stem cell apoptosis. Specifically, C5aR1 signalling attenuates a macrophage maturation response characterised by increased IL10 and CX3CR1 expression. Our data indicate that increased IL10 signalling, and macrophage maturation induced following C5aR1 targeting are important for regulating crypt cell loss via radiotherapy-induced apoptosis. Interestingly, IL10-signalling is also critical for the improved tumour radiation response observed following C5aR1 targeting.

## Introduction

In the intestine, stem cells located at the bottom of the crypt are responsible for intestinal homeostasis and regeneration^1^. Injury, which may occur following cytotoxic cancer treatment, compromises intestinal integrity and function when regenerative stem cells in the intestinal crypt are lost following apoptosis and regeneration programmes are not efficiently deployed^2,3^. Macrophages have emerged as key for regulating stem cell regeneration following injury^4–6^. However, how to therapeutically promote specific macrophage phenotypes with the capacity to tip the balance towards regeneration and away from apoptosis is still unclear. Identification of key molecular regulators of macrophage phenotypes in the intestinal niche, as well as how these are affected by treatment, is essential for understanding how intestinal function is maintained during homeostasis and injury^3^. This understanding will be critical for identifying novel targets to reduce toxicity from inflammatory cytotoxic cancer treatments while retaining effective tumour control.

Approximately 300,000 patients worldwide receive pelvic irradiation every year and 50-70% of these patients experience acute toxicity^7–9^. In fact, the small intestine is often subject to some of the most distressing damage when incidentally present in the field during pelvic, upper gastrointestinal tract, inferior lung or retroperitoneal irradiation^9,10^. The resulting condition is known as radiation-induced small bowel disease (also historically known as radiation enteritis or enteropathy)^9^. In fact, in clinical practice, minimising toxicity associated with radiotherapy (RT) while still achieving maximal tumour control (i.e. achieving the most optimal therapeutic window) is actively considered when preparing treatment plans^11–13^. Opening the therapeutic window of RT through normal tissue radioprotection would be a critical step in RT dose escalation, raising the possibility of using RT with curative intent in an increasing number of settings. Despite the importance of considering both tumour and normal tissue responses in clinical practice, few preclinical studies have comprehensively investigated the effects of RT-modulating therapies on both tumour and normal tissue responses^14^. As a consequence, there is currently no approved agent that can both radiosensitise tumours and radioprotect normal tissue. We recently described C5aR1 as an innate immunity target that can improve tumour RT response, including in tumours with immunosuppressive microenvironments associated with the worse prognostic outcome^15^. Here we provide evidence that C5aR1 targeting can be radioprotective of the normal intestine. We identify that increased IL10 signalling and macrophage maturation are key for radiation-induced apoptosis regulation in the intestinal crypts, likely supporting an expedited damage resolution response following C5aR1 targeting. Interestingly, IL10 signalling is also critical for the improve tumour radiation response observed following C5aR1 targeting.

## Results

### Targeting C5aR1 reduces histological damage and improves survival following RT

We recently reported that targeting C5aR1 can improve tumour response through increasing tumour cell apoptosis but without resulting in increased apoptosis in the healthy intestinal crypt^15^. However, how C5aR1 targeting impacts radiation-induced small bowel disease/ intestinal injury has not been comprehensively evaluated. To this end, we first established a histologic scoring system, whereby increasing levels of RT-induced intestinal injury were assessed through the scoring of 6 different histologic parameters in mice **(Supplementary Table 1 and Figure 1A and B).** Briefly, histologic parameters included neutrophilic inflammation (and distribution), crypt damage (and distribution), and crypt regeneration impairment (and distribution). All slides were blindly evaluated by a board-certified veterinary pathologist at two-days post-RT, a timepoint at the interphase between the apoptotic and regenerative/proliferative response to RT. We confirmed the validity of our damage scoring system by irradiating mice with increasing doses of total abdominal irradiation. As shown in **Figure 1B**, increasing RT doses resulted in corresponding increases in total damage scores within the small intestine.

**Figure 1.**
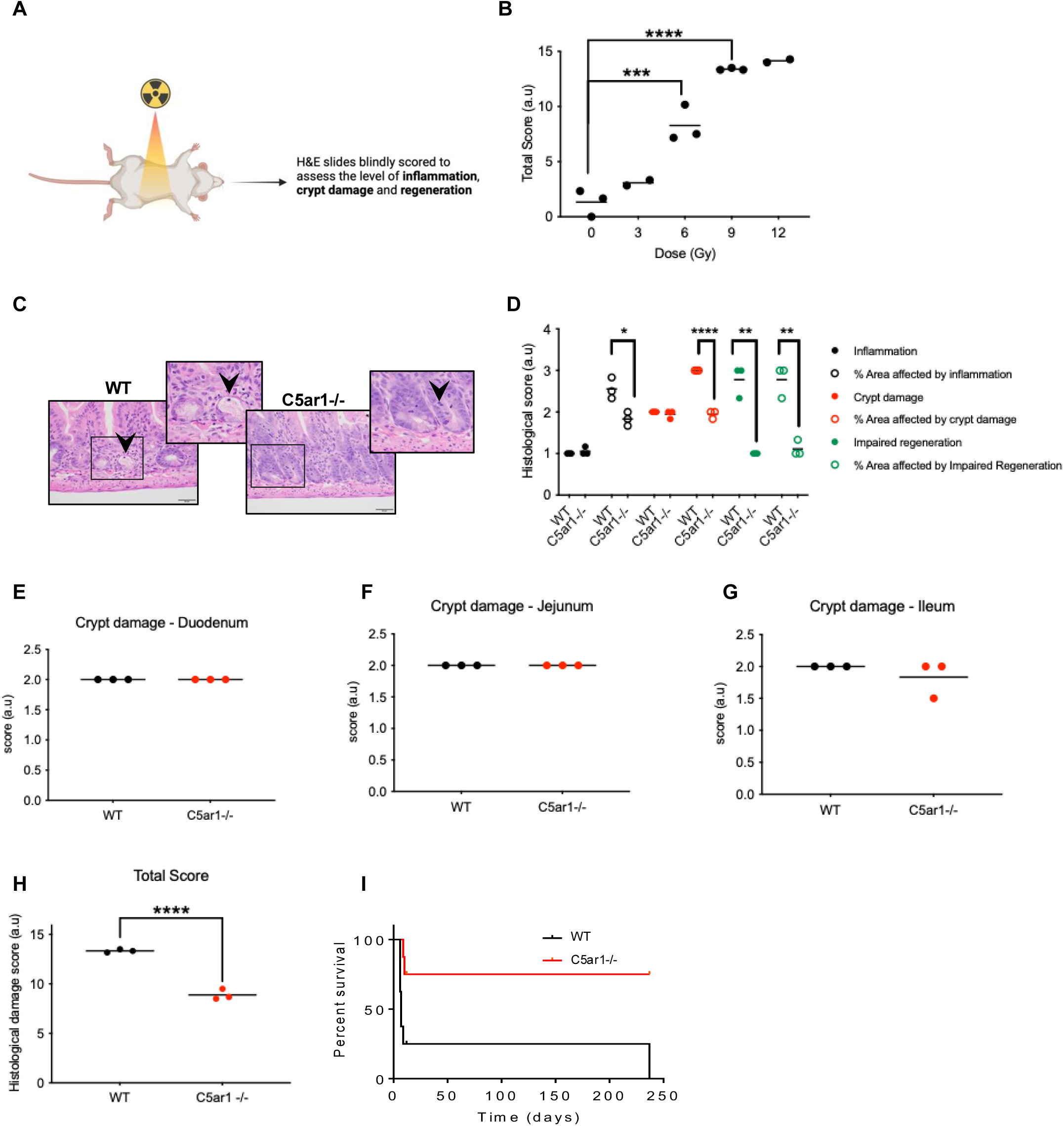
Targeting C5aR1 reduces histological damage and improves survival following RT. **(A)** Schematic representation of experimental design. **(B)** Graph shows an increase in the total histologic damage score with increasing irradiation dose. Total histologic damage score obtained by adding scores of 6 individual histologic parameters. Intestines were harvested 2 days post-RT. See also Supplementary Table 1. **** = p<0.0001, *** = p<0.001 by 1 way ANOVA with Dunnett’s comparison test. Individual points represent individual mice per group. **(C)** Representative image of H&E staining in WT and C5ar1^-/-^ mice. **(D)** Graph shows the breakdown of individual scores used to calculate total histological damage scores for WT and C5ar1^-/-^ mice irradiated with 9 Gy total abdominal irradiation. Intestines were harvested 2 days post-RT. Slides scored blindly by a pathologist. **** = p<0.0001, ** = p<0.01 2-tailed t-test. Individual points represent individual mice per group. **(E)** Graph shows the crypt damage score in the duodenum of BALB/c mice following 9 Gy total abdominal irradiation. Intestines were harvested 2 days post-RT. Individual points represent individual mice per group. **(F)** Graph shows the crypt damage score in the jejunum of BALB/c mice following 9 Gy total abdominal irradiation. Intestines were harvested 2 days post-RT. Individual points represent individual mice per group. **(G)** Graph shows the crypt damage score in the ileum of BALB/c mice following 9 Gy total abdominal irradiation. Intestines were harvested 2 days post-RT. Individual points represent individual mice per group. **(H)** Graph shows total histologic damage score of WT and C5ar1^-/-^ mice. Intestines were harvested 2 days post-RT. **** = p<0.0001, two tailed t-test. Individual points represent individual mice per group. **(I)** Kaplan-Meier curve for WT or C5ar1^-/-^ mice irradiated with 12 Gy total abdominal irradiation. * = p<0.05, 2-tailed t-test, n = 7 per group.

We used our scoring system as a benchmark to compare the effects of C5aR1 loss on RT-induced toxicity. We observed that C5ar1^-/-^ mice displayed a significantly decreased damage score relative to WT mice which was evident even when all six parameters were assessed separately **(Figure 1C, D and Supplementary Figure 1A)**. Of note, the total crypt damage score was the same between the two groups of mice, consistent with the fact that both groups received the same irradiation dose **(Figure 1D)**. Also, the same scores were observed along the length of the small intestine, with equivalent crypt damage, observed in the duodenum, jejunum and ileum **(Figures 1E, F and G).** As anticipated from analysis of separate inflammation, crypt damage and regeneration impairment scores, assessment of the total levels of histological damage indicated that C5ar1^-/-^ displayed a significantly decreased total score relative to WT mice **(Figure 1H)**.

To assess the longer-term effect of C5aR1 loss on the intestinal response to RT, we compared survival in WT and C5ar1^-/-^ mice following a dose of 12 Gy (expected to kill >90% of WT mice in this strain). We found that long-term survival was significantly improved in C5ar1^-/-^ compared to WT mice **(Figures 1I)**. These data indicate that C5aR1 depletion in mice results in decreased small intestinal histologic damage score and increased survival following abdominal irradiation.

### Macrophage maturation and protective radiation responses

To investigate the mechanisms underlying the radioprotective effects, spatial transcriptomics (GeoMx) of irradiated intestinal crypt regions of WT and C5ar1^-/-^ mice was undertaken **(Figure 2A)**. Analysis of pathways differentially expressed between WT and C5ar1^-/-^ mice indicated that, as expected, G-Protein Coupled Receptor (GPCR) signalling was downregulated in C5ar1^-/-^ compared to WT mice **(Figure 2A and Supplementary Table 2)**.

**Figure 2.**
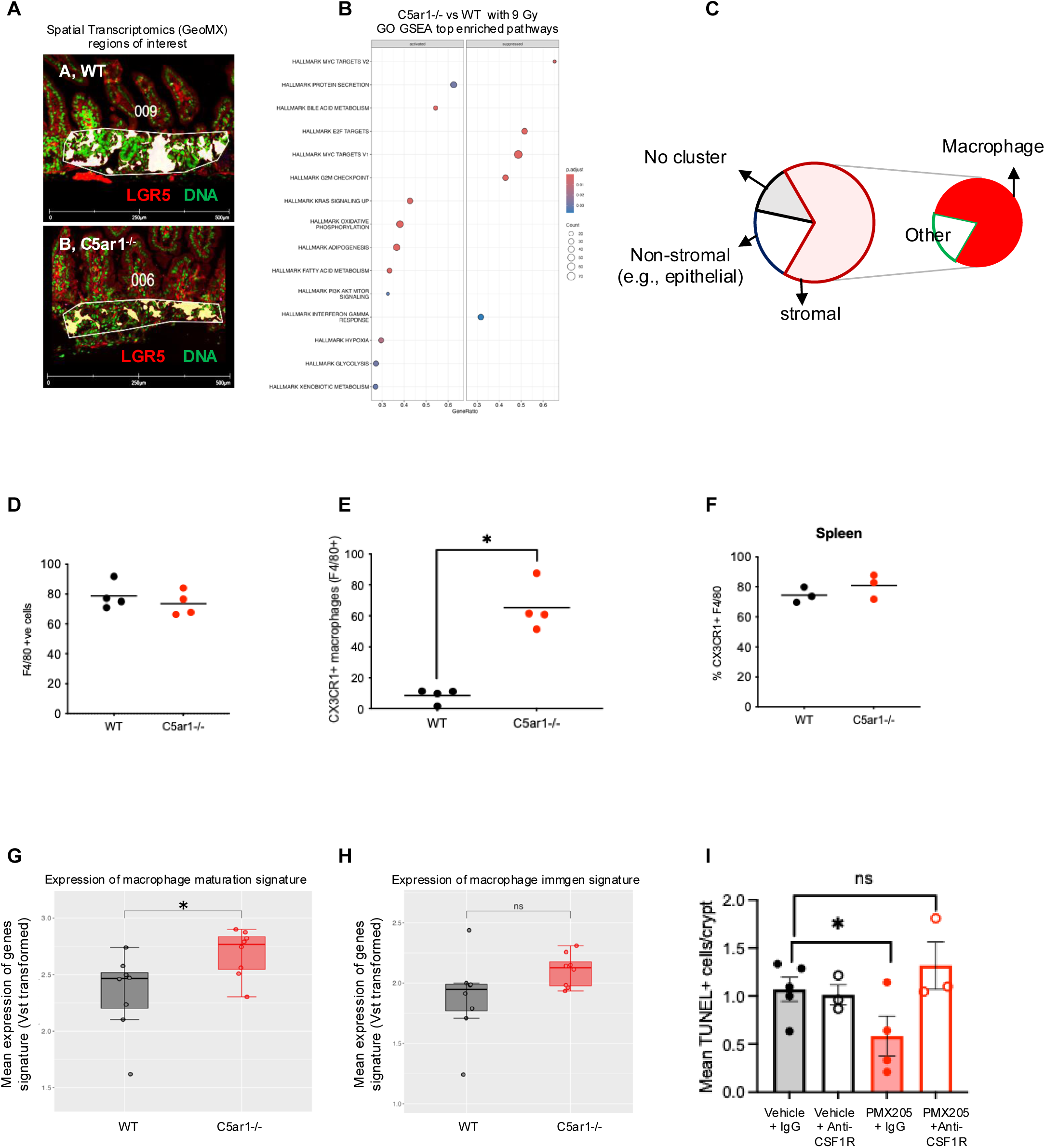
Macrophage maturation and protective radiation responses. **(A)** Representative image of spatial transcriptomics regions of interest. **(B)** Dot plot shows top enriched pathways following GSEA of spatial transcriptomics data. Intestinal crypt regions of WT and C5ar1^-/-^ mice (treated with 9 Gy total abdominal irradiation) were harvested 3 days post-RT. **(C)** Pie chart representing cells primarily expressing C5aR1-regulated Genes of Interest (GOIs) as identified following GSEA of spatial transcriptomics data from A. **(D)** Graph shows the % of F4/80+ cells found in small intestine of WT or C5ar1^-/-^ mice. Individual points represent individual mice per group. **(E)** Graph shows the % of CX3CR1+ macrophages found in small intestines of WT or C5ar1^-/-^ mice. * = p<0.05, 2-tailed t-test. Individual points represent individual mice per group. **(F)** Graph shows the % of CX3CR1^+^ F4/80 macrophages found in spleens of WT or C5ar1^-/-^ mice. Individual points represent individual mice per group. **(G)** Expression of macrophage maturation signature (from spatial transcriptomics data) in WT and C5ar1^-/-^ mice following 9 Gy irradiation. Intestines were harvested 3 days post-RT. **(H)** Expression of Immgen macrophage signature (from spatial transcriptomics data) in WT and C5ar1^-/-^ mice following 9 Gy irradiation. Intestines were harvested 3 days post-RT. **(I)** Graph shows the number of TUNEL + cells in C57/BL6 mice irradiated with 9 Gy total abdominal irradiation and treated with either IgG or anti-CSF1R antibody. Intestines were harvested 2 days post-RT. * = p<0.05, ** = p<0.01, 2-tailed t-test. Individual points represent individual mice per group.

Gene set enrichment analysis (GSEA) indicated that the most significantly downregulated genes in C5ar1^-/-^ mice were associated with cell cycle/proliferation (e.g., G2/M checkpoint, E2F targets, **Figure 2B**). However, no significant difference was observed in the staining of proliferation marker Ki67 in tissues from mice with either genetic or pharmacological targeting of C5aR1 following RT treatment (**Supplementary Figures 2A-C**). These data suggest that changes in proliferative signalling may not be driving the radioprotective response observed following C5aR1 targeting.

Further inspection of differentially expressed genes between C5ar1^-/-^ and WT mice indicated that the majority of these were associated with expression by stromal populations rather than epithelial cells **(Figure 2C)**. Notably, from the top 10 differentially expressed genes, 40% were genes known to be expressed primarily by stromal rather than epithelial cells and within this group, 80% were associated with macrophage/myeloid expression **(Figure 2C and Supplementary Table 3)**. Using mMCP-counter and SeqImmuCC spatial deconvolution algorithms we inferred the immune cell populations present in the sequenced crypt regions (**Supplementary Figures 2D and E)**. Macrophages were the most abundant stromal populations identified by both algorithms with mMCP-counter identifying a significant difference in monocytes in the C5ar1^-/-^ mice. SeqImuCC estimated a lower % of CD8^+^ T-cells in C5ar1^-/-^ mice, a finding validated by flow cytometry analysis indicating that, at baseline, C5ar1^-/-^ mice already displayed reduced intestinal CD8^+^ T cells and increased numbers of intestinal CX3CR1^+/^F4/80^+^ macrophages despite total numbers of CD45^+^ cells and F4/80^+^ macrophages being comparable between both groups of mice **(Figure 2D, 2E and Supplementary Figures 2F-H)**. No changes in Cx3cr1^+/^F4/80^+^ macrophages were observed in the spleen suggesting this is a local intestinal-tissue specific effect **(Figure 2F)**. These data are in agreement with a lack of changes observed in the global macrophage immgen transcriptional signature in the crypts of C5ar1^-/-^ mice which, however, did display significantly increased expression of genes associated with macrophage maturation (which include *Cx3cr1*) following RT (**Figures 2G and 2H)**. Since macrophages are important for apoptotic cell clearance and we have previously reported reduced apoptosis in intestinal crypts following pharmacological or genetic C5aR1 targeting, we investigated whether an altered macrophage maturation phenotype following C5aR1 targeting could support apoptosis clearance^15^. CSF1-CSF1R signalling is essential for monocyte and macrophage proliferation and maturation^16^. We found that CSF1R blockade, (which resulted in a 10-15% reduction in intestinal macrophages), did not significantly alter levels of apoptosis in vehicle treated mice following RT **(Figure 2I and Supplementary Figure 2I)**. However, the reduced apoptosis previously observed following C5aR1 targeting was abrogated upon CSF1R blockade **(Figure 2I and Supplementary Figure 2I).** These data highlight the importance of macrophages and their maturation for reduced crypt cell apoptosis observed following C5aR1 targeting.

### C5aR1 signalling prevents apoptotic cell clearance through an attenuated IL10-dependent response

To gain further insights into the role of different macrophage subsets in RT-induced intestinal injury, we interrogated single-cell RNA-sequencing (scRNA-Seq) data of small intestines before and after abdominal RT **(Figure 3A)**. We divided macrophages into 4 groups (muscularis, mucosal, mature and intestinal monocytes) and compared expression of genes previously associated with inflammatory resolution and repair (*Mrc1/cd206*, *Cd163*, *Il10* and *Cx3cr1*) before and after RT. Interestingly, we found that mature macrophages were the only group that expressed all 4 genes before RT, with *Il10* being expressed at the lowest levels in relation to the other genes **(Figure 3B)**. Interestingly, following RT, *Cx3cr1* and *Il10* expression in this subset appeared abrogated compared to the other two genes suggesting that irradiation may preferentially affect expression of these markers **(Figure 3C)**.

**Figure 3.**
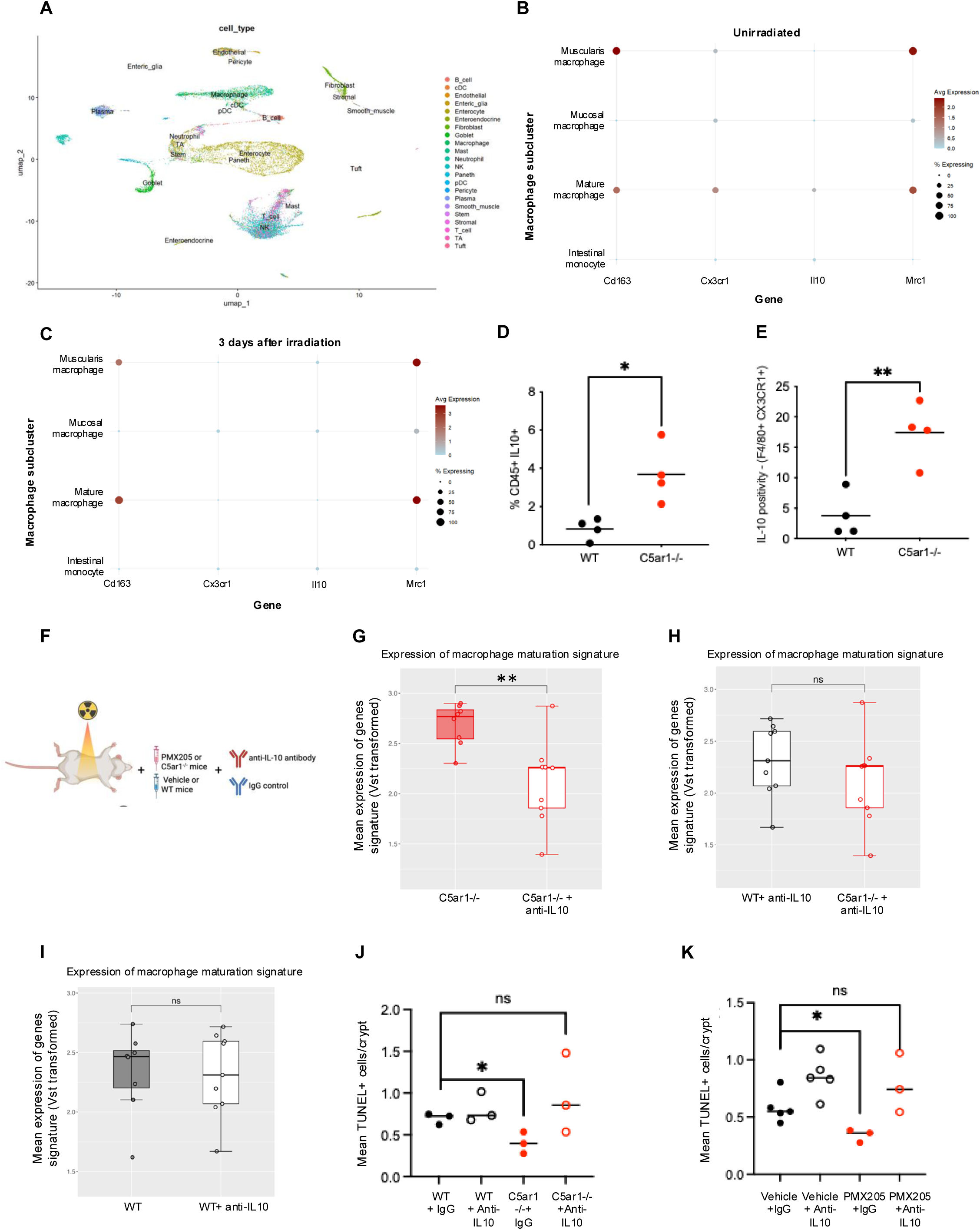
C5aR1 signalling prevents apoptotic cell clearance through an attenuated IL10-dependent response. **(A)** UMAP of 17, 693 cells distributed by annotated unsupervised clustering. **(B)** Dot plot showing expression of specific macrophage-associated genes in different macrophage subtypes in unirradiated mouse small intestines. Mean expression reflects expression relative to other cell types and genes in the graph. Expression levels are log-normalised and scaled. The percent expressing reflects the proportion of cells of each type expressing the gene. **(C)** Dot plot showing expression of specific macrophage-associated genes in different macrophage subtypes in irradiated mouse small intestines 3 days after RT. Mean expression reflects expression relative to other cell types and genes in the graph. Expression levels are log-normalised and scaled. The percent expressing reflects the proportion of cells of each type expressing the gene. **(D)** Graph shows the % of CD45^+^ and IL-10^+^ cells found in small intestines of WT or C5ar1^-/-^ mice. * = p<0.05, 2-tailed t-test. Individual points represent individual mice per group. **(E)** Graph shows the % IL-10 positivity in F4/80^+^ and CX3CR1^+^ cells found in small intestines of WT or C5aR1^-/-^ mice. ** = p<0.01, 2-tailed t-test. Individual points represent individual mice per group. **(F)** Schematic representation of experimental design. **(G)-(I).** Expression of macrophage maturation signature (from spatial transcriptomics data including from Figure 2A) in WT and C5ar1^-/-^ mice receiving anti-IL-10 antibody or IgG2b treatment and 9 Gy total abdominal irradiation. Intestines were harvested 3 days post-RT. **(J)** Graph shows the number of TUNEL+ cells in WT or C5ar1^-/-^ mice irradiated with 9 Gy total abdominal irradiation and treated with 3 doses of either IgG or IL-10 blocking antibody (flanking the irradiation). Intestines were harvested 3 days post-RT. * = p<0.05, 2-tailed t-test. Individual points represent individual mice per group. **(K)** Graph shows the number of TUNEL + cells in mice irradiated with 9 Gy total abdominal irradiation and treated with 3 doses of either IgG or IL-10 blocking antibody (flanking the irradiation) +/- PMX205. Intestines were harvested 3 days post-RT. * = p<0.05, 2-tailed t-test. Individual points represent individual mice per group.

IL10 can be produced by both epithelial as well as immune (CD45^+^) cells^17,18^. We found that both CD45^-^ and CD45^+^ cells in the small intestines of WT and C5ar1^-/-^ mice expressed IL10 **(Figure 3D and Supplementary Figure 3A),** with C5ar1^-/-^ mice displaying significantly increased IL10 levels only in CD45^+^ cells **(Figure 3D)**. Within the CD45^+^ population, C5ar1^-/-^mice displayed increased levels of F4/80^+^ CX3CR1^+^ macrophages expressing IL10 in the intestine but not in the spleen **(Figures 3E and Supplementary 3B).** CD4^+^ cells in the intestines of C5ar1^-/-^ mice also expressed increased IL10, albeit this did not reach statistical significance **(Supplementary Figure 3C)**.

To assess whether IL10 signalling and macrophage maturation phenotypes are important for the protection phenotype observed following C5aR1 targeting, we used IL10 blocking antibodies in mice following either pharmacological or genetic C5aR1 targeting **(Figure 3F**). Following analysis of spatial transcriptomics data (including from Figure 2), we noted that the general increase in macrophage maturation signatures observed in the crypts of C5ar1^-/-^mice was reversed upon IL10 blockade **(Figure 3G-I)**. Importantly, in line with the changes in macrophage maturation phenotype, mice treated with IL10 blockade in combination with C5aR1 targeting (pharmacological or genetic) had increased levels of crypt cell apoptosis compared to mice where C5aR1 targeting occurred in combination with IgG control antibodies **(Figure 3J and 3K**). Together these data indicate that targeting C5aR1 productively engages the IL-10-macrophage axis to favour reduced crypt cell apoptosis following RT.

### Increased radiosensitivity following C5aR1 targeting is IL10-dependent

While reduced apoptosis following C5aR1 targeting is beneficial in the context of normal tissue injury, reduced IL10-dependent tumour cell death could limit tumour RT responses. We therefore assessed the effects of IL10 blockade on colorectal cancer cell death/apoptosis following RT and treatment with or without C5aR1 targeting. In agreement with our previous studies^15^, C5aR1 antagonist (PMX205) and RT treatment in human colorectal cancer cells (HT29 and HCT116) *in vitro*, resulted in increased apoptosis/cell death compared to cells treated with RT alone **(Figures 4A and B)**^15^. This effect was also observed following TUNEL staining of sections from colorectal MC38 syngeneic tumour xenografts treated with PMX205 and RT **(Figures 4C and D)**. Interestingly, IL10 blockade in combination with C5aR1 targeting reduced cell death/apoptosis both *in vitro*, and in MC38 syngeneic tumour models *in vivo* **(Figures 4A-D)**. To more directly assess the effect of IL10 blockade on tumour responses, we treated MC38 tumour xenografts with IL10 blocking antibodies together with RT and PMX205 and assessed tumour growth delay following RT. Consistent with the *in vitro* and TUNEL staining results, PMX205 treated tumours displayed improved tumour RT response compared to vehicle treated tumours but this effect was abrogated upon IL10 blockade **(Figure 4E and F)**. These data indicate that following C5aR1 targeting, IL10 plays a key role in both normal tissue radioprotection and tumour radiosensitisation.

**Figure 4.**
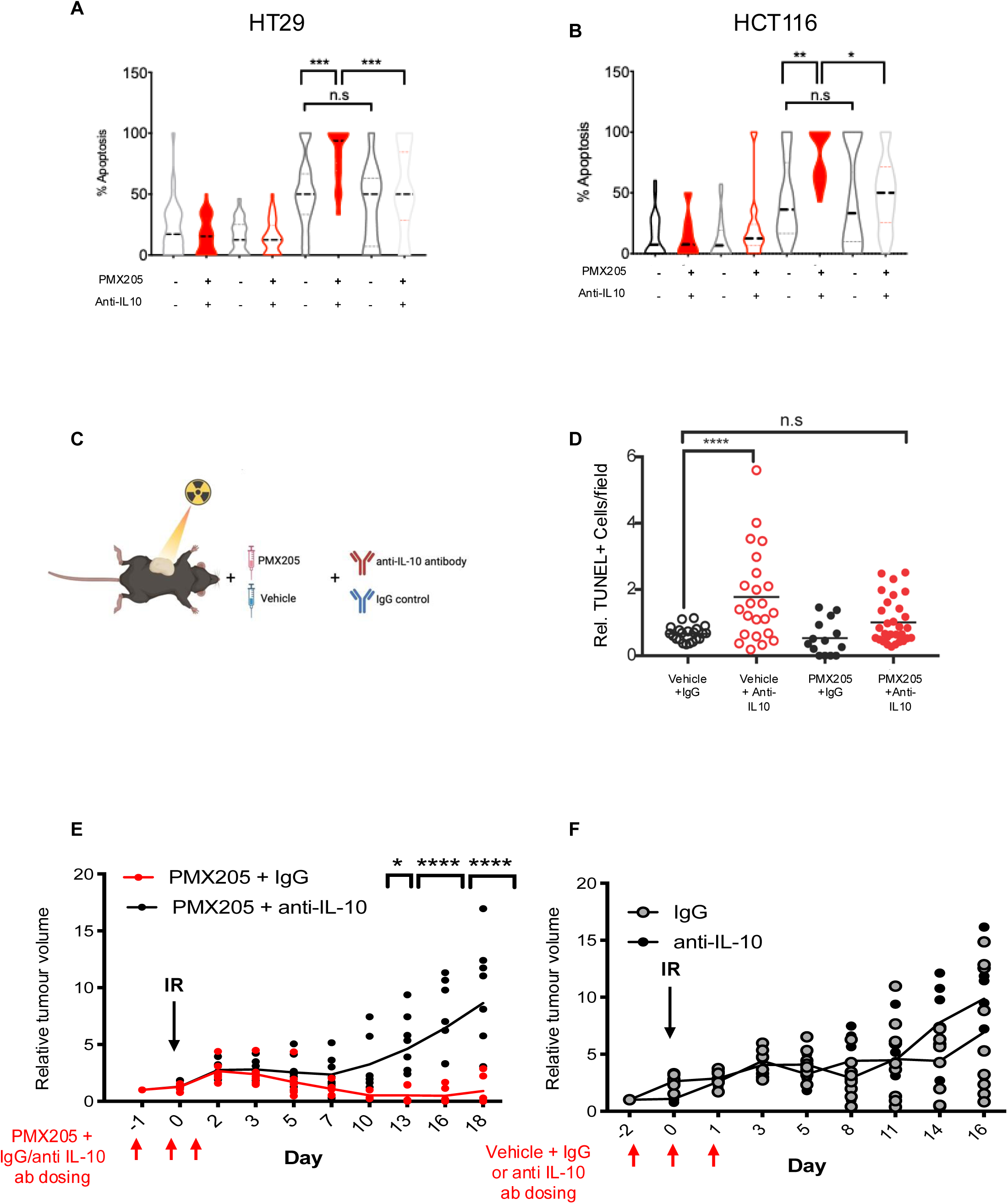
Increased radiosensitivity following C5aR1 targeting is IL10-dependent. **(A)** The graph represents the number of apoptotic/non-apoptotic cells expressed as a % of the whole population for HCT116 cells treated with either vehicle or PMX205 and either IgG or IL-10 blocking antibody from 1 hour before irradiation with either 0 or 9 Gy. Cells were harvested 48 hours post-RT. n=3. **** = p<0.0001 by 2-way ANOVA with Dunnett’s comparison test. **(B)** The graph represents the number of apoptotic/non-apoptotic cells expressed as a % of the whole population for HT29 cells treated with either vehicle or PMX205 and either IgG or IL-10 blocking antibody from 1 hour before irradiation with either 0 or 9 Gy. Cells were harvested 48 hours post-RT. Independent fields of view are show, n=3. **** = p<0.0001 by 2-way ANOVA with Tukey-Kramer comparison test. **(C)** Schematic representation of experimental design. **(D)** The graph represents the number of TUNEL+ cells found in MC38 subcutaneous tumours treated with either vehicle or PMX205 and either IgG or IL-10 blocking antibody 1 day before irradiation (10 Gy), on the day of and 1 day post-irradiation. Individual points represent independent fields of view from up to 3 different tumours per group. **** = p<0.0001, n.s = not significant, 2-way ANOVA with Dunnett’s comparison test. **(E)** Relative tumour growth curves are shown for MC38 colorectal cancer cells treated with 10 Gy single dose irradiation, PMX205 treatment and either IgG or IL-10 blocking antibody for 3 doses flanking the irradiation dose (on day 0, 1 and 2). * = p<0.05, ** = p<0.01 **** = p<0.0001 by 2-way ANOVA. Individual points represent individual mice per group. **(F)** Relative tumour growth curves are shown for MC38 colorectal cancer cells treated with 10 Gy single dose irradiation, vehicle treatment and either IgG or IL-10 blocking antibody for 3 doses flanking the irradiation dose (on day 0, 1 and 2). Individual points represent individual mice per group.

## Discussion

As a first line of defence against foreign substances, the innate immune system senses pathogen- or danger-associated molecular patterns and initiates an inflammatory response to clear the danger^19^. In infection, this initial inflammatory burst is important for containing infection, but this response must be eventually contracted to prevent tissue damage. For example, production of IL10 by phagocytes is critical for ensuring a return to cellular homeostasis following inflammation thereby preventing autoimmunity^20^. In the immunotherapy era, inducing an enhanced inflammatory response is emerging as a popular strategy to promote anti-tumour immunity and tumour clearance^21,22^. However, failure to contract this inflammatory burst in a timely manner during standard-of-care cytotoxic cancer treatment could result in normal tissue toxicity and detrimental side effects impacting patients’ quality of life^23^. How inflammation resolution is coordinated with apoptosis clearance and regeneration attempts following inflammatory cytotoxic cancer treatments such as RT has remained unclear.

Here we uncover C5aR1 as a novel target to reduce RT-induced intestinal injury. Mechanistically, C5aR1 loss results in increased abundance of CX3CR1^+^/IL10^+^ macrophages in the intestine and enhanced IL10-signalling modulates intestinal macrophage maturation phenotypes relevant for the radioprotection phenotype observed following C5aR1 targeting. In particular, the reduced intestinal crypt cell apoptosis observed following C5aR1 targeting is CSF1R- and IL10-dependent. CSF1R neutralisation has been previously shown to preferentially impair maturation (and number) of tissue resident macrophages, without having a major impact on monocytes^24^. The absence in changes in apoptosis levels following CSF1R blockade in vehicle (rather than PMX205 treated mice) could indicate a higher dependence on tissue resident macrophages in the PMX205 treated group.

Interestingly, IL10 signalling in high CX3CR1-expressing macrophages polarises them to an anti-inflammatory phenotype that may further contribute to the reduced RT-induced injury observed^17,18^. Indeed, during tissue injury, macrophages are involved in all stages of repair, and after initially regulating inflammation, they redirect the inflammatory response towards repair, regeneration and finally injury resolution^25^. It is tempting to speculate that the radioprotection observed following C5aR1 targeting is underlined by a skewing of macrophage populations towards this anti-inflammatory pro-resolving state that primes the tissue to have a more effective injury repair response.

Interestingly, our data indicate that IL10 mediates the opposite apoptosis effects in the small intestine and in tumours. Namely, apoptosis is increased in an IL10-dependent manner following C5aR1 targeting in tumours while it decreases in intestinal crypts. We hypothesize that increased IL10 poises tumours to respond to RT through increased apoptosis propensity. This may occur via IL-10 dependent downregulation of NF-κB pro-survival signalling which we have previously show is important for tumour radiosensitisation following C5aR1 targeting^15,25^. However, this remains to be formally proven.

Overall, our data indicate that C5aR1 is a promising therapeutic target for improving both tumour response and normal tissue radioprotection with increased IL10 signalling underlying the improved tumour and normal tissue responses observed.

## Materials and Methods

### Cell lines and treatments

HCT116 male adult human epithelial colorectal carcinoma and HT-29 female adult colorectal adenocarcinoma cells were purchased from ATCC**^®^.** MC38 murine C57BL6 colon adenocarcinoma cells were provided in-house. Cells were grown in DMEM with 10% FBS, in a standard humidified incubator at 37°C and 5% CO_2_. All cell lines were routinely tested for mycoplasma and found to be negative.

### IL-10 blocking antibody treatment

For all *in vitro* experiments, cells were pre-treated with 1 μg/ml IL-10 antibody (R&D # MAB217-100) 1 hour before irradiation. Mouse anti-human IgG (Santa Cruz, #sc-2025) was used as the IgG control.

For *in vivo* experiments, IL-10 blocking antibody (LEAF Purified clone JES5-16E3, Biolegend #505012)(50 μ_g_ 24 hours before irradiation, 100 μ_g_ on the day of irradiation and 15 μ_g_ 24 hours post irradiation) or IgG control (LEAF Purified rat IgG2b, κ isotype control #400637, Biolegend) (same dosing and concentration as IL-10 antibody) were used.

### CSF1R blocking antibody treatment

Mice were treated with three doses (200 μg) of either BioXCell InVivoMAb anti-mouse CSF1R (BE0213) or BioXCell InVivoMAb rat IgG2a isotype control, anti-trinitrophenol (BE0089). Doses were given 48 hours before RT (9 Gy), 24 hours before RT and on the day of RT.

### PMX205 treatment

For *in vitro* experiments, cells were treated with 10 μg/ml PMX205 (Tocris #5196) dissolved in 20% ethanol/water. Vehicle control in these experiments refers to 20% ethanol/water. Cells were pre-treated for 1 hour before irradiation.

For *in vivo* experiments involving PMX205 treatment, 10 mg/kg PMX205 (Tocris #5196, or synthesized and purified as previously described^26^) was administered to mice (orally) flanking the irradiation doses. Vehicle control in these experiments refers to 20% ethanol/water.

### Animal studies

Experimental procedures were carried out under a project license issued by the UK Home Office under the UK Animals (Scientific Procedures) Act of 1986; or in accordance with the NIH guidelines for the use and care of live animals (approved by the Stanford University Institutional Animal Care and Use Committee). Animal Research Reporting of In vivo Experiments (ARRIVE) guidelines were used.

For all studies, animals were randomly divided into groups using a computer-based random order generator and blinding during data collection and analysis was carried out whenever possible.

For normal tissue toxicity studies, total abdominal irradiations (except for those experiments shown in in Figure 3K) were performed on anaesthetised animals (with the use of ketamine 100 mg/kg/xylazine 20mg/kg) and using a 225 kVp cabinet X-ray system filtered with 0.5 mm Cu (at Comparative Medicine Unit, Stanford, CA). The Gulmay Medical RS320 irradiator (300 kV, 10 mA, 1.81 Gy/min with isoflurane anaesthesia, 3% isoflurane and 100% O_2_ for induction, and 2.5% isoflurane for maintenance) was used for RT for experiments shown in Figure 3K.

To perform irradiations of subcutaneous tumours, The X-Rad SmART (Precision X-Ray Inc., North Branford, CT) was used as in^15^. Mice were anaesthetised in a knockdown chamber with a mixture of 3% isoflurane and 100% O_2_. Anaesthesia was maintained using 1.5% isoflurane in O_2_ delivered via a nose cone. Mice were monitored for breathing with a webcam fitted in the cabinet. The prescribed dose was calculated for a point chosen approximately in the middle of the tumour. Pre-treatment computed tomography (CT) images were acquired to facilitate treatment planning, using a beam energy of 40 kVp, a beam filter of 2 mm Al, and a voxel size of 0.2 or 0.1 mm. CT images collected were loaded onto RT_Image and a treatment plan was created using a single beam (using open-source RT_Image software package, version 3.13.1, running on IDL version 8.5.1^27^). Therapeutic irradiations were performed using an x-ray energy of 225 kVp, a current of 13 mA, a power of 3000 Watts, and a beam filter of 0.3 mm Cu, producing a dose rate of ∼300 cGy/min at the isocenter. Treatment x-ray beams were shaped using a 10 or 15 mm collimator to selectively irradiate the target while sparing adjacent tissue when performing xenograft irradiation.

For IL-10 neutralisation experiments, MC38 cells (5 x 10^5^) were injected subcutaneously into 6-8 week old female C57/BL6 (JAXX) at a single dorsal site. Growing tumours (average, 80-100 mm^3^) were treated with 10 Gy (as described above).

### Histology

Following euthanasia, small intestinal tissues were harvested and immersion-fixed in 10% neutral buffered formalin for 72 hours. Following fixation, serial longitudinal sections of small intestine were submitted for routine sectioning (at Histo-Tec Laboratory, Inc. Hayward, CA or in house). Briefly, slides were routinely processed, embedded in paraffin, section at 5 μm, and stained with hematoxylin and eosin (HE). To assess radiation-induced toxicity, an ordinal grading scale was created that evaluated the following six parameters: 1) neutrophilic inflammation, 2) tissue area (%) affected by neutrophilic inflammation, 3) crypt damage, 4) tissue area (%) affected by crypt damage, 5) crypt regeneration impairment, and 6) tissue area (%) affected by crypt regeneration impairment (see Supplementary Table 1 for scoring definitions). Six sections of small intestine (duodenum, jejunum, and ileum) were evaluated per mouse. A total damage score was calculated for each mouse by adding individual scores across all six individually evaluated parameters (highest score possible = 18). All sections were blindly evaluated by a single board-certified veterinary pathologist (K.M.C.).

## Immunohistochemistry

For Ki67 staining shown in Supplementary Figures 2A and B: formalin-fixed paraffin-embedded sections were dewaxed by soaking the slides in xylene for 15 minutes followed by 5 minute washes in 100%, 90%, 80%, and 70% ethanol, and 5 min in 3% hydrogen peroxide. The sections were then placed in citrate buffer (10mM citric acid, 0.05% Tween-20) and microwaved for 10 minutes for antigen retrieval. Quenching was performed with 1% hydrogen peroxide in methanol for 20 min. Sections were stained for Ki67 (Themo Fischer Scientific MA5-14520). After incubation with secondary antibody—biotinylated anti-horse IgG (Vector Laboratories, BA-8000) and biotinylated anti-goat IgG (Vector Laboratories, BA-9500), avidin-biotin complex technique was performed with the ABC kit (Thermofisher, 32020) to enhance detection, which was accomplished using the DAB Substrate Kit (Abcam, ab64238) for 45 seconds. Counterstaining of nuclei was done with hematoxylin for 30 seconds then dehydrated with ethanol. Slides were mounted using DPX mounting medium and imaged. Stained slides were scanned with a NanoZoomer 2.0-RS Digital Slide Scanner (Hamamatsu). The two sections were overlaid and 250 µm x 250 µm squares were magnified individually. Image saturation was increased to 200% to increase contrast between the blue-stained nuclei and the brown-stained marker of interest. Staining was quantified by counting number of positive cells/crypts in 10 random high-power fields per slide.

For the immunofluorescent Ki67 staining shown in Supplementary Figures 2C: 5 µm FFPE sections were incubated at 50 °C for 10 min. Two, 5-min Citroclear incubations at room temperature deparaffinised the tissues. Tissues were rehydrated by 100%, 100%, 70% and 50% ethanol incubations for 5 min at room temperature. For antigen retrieval, slides were incubated at 100 °C and 5 psi for 5 min in a container of 100 mL of 0.05% Tween 20 in 1 X citrate buffer. The slides were cooled to room temperature and washed with 0.05% Tween 20 in PBS (PBS-T) on a shaker for 3 min. Non-specific antibody binding was limited by blocking with Protein Block Serum-Free Ready-to-use solution (DAKO, X0909) for 30 min at room temperature. The slides were washed for 3 min with PBS-T. Sections were incubated overnight at 4 °C in 1: 100 primary rabbit anti-Ki67 antibody (Abcam, AB16667) diluted in PBS-T. Excess antibodies were removed by two 3-min PBS-T washes. Sections were incubated at room temperature for 90 min in 1: 500 donkey anti-rabbit IgG (H+L) highly cross-absorbed secondary antibody, Alexa Fluor™ Plus 594 (Invitrogen, A32754) diluted in Protein Block. Two 3-min PBS-T washes were performed before mounting the slides with ProLond Gold Antifade Mountant with DNA Stain DAPI (Invitrogen, P36931). The slides were imaged using a MICA microscope (Leica Microsystems).

### TUNEL assay

ApopTag® (Millipore #S7100) was used to stain 4 mm formalin-fixed paraffin-embedded small intestinal or tumour sections. Stained slides were scanned with a NanoZoomer 2.0-RS Digital Slide Scanner (Hamamatsu) or Aperio CS scanner (Aperio Technologies). For tumour studies, fifteen randomly chosen fields of view at 10x magnification were selected for analysis via ImageJ software (NIH). Images were first converted to 8-bit black and white, then the threshold was adjusted to 0-200 to eliminate background signal. The remaining particles larger than 10 square pixels were counted. The % of counts/total area analysed per field is shown. For small intestines, the number of TUNEL+ cells per crypt were manually counted in at least 10 fields of view per section.

Apoptosis assessment by morphology in cell lines was carried out as previously described ^28,29^. Adherent and detached cells (in media) were collected and fixed in 4% PFA for 15 minutes at room temperature. PFA was then removed, and cells were washed in 1 ml PBS. 10 μl of fixed cells in PBS was placed in each slide and mixed with ProLong^TM^ Gold antifade mountant with DAPI (Invitrogen #P36935). A coverslip was placed on top of the sample and slides were left to dry overnight. Slides were imaged using a DSM6000, DMi8 or DMI6000 (Leica) microscope with 40x or 60x oil objectives. The number of cells with fragmented DNA and the total number of cells per field was counted (with typically at least 10 fields counted for every treatment).

### Survival of mice after irradiation

Actuarial survival was calculated by the Kaplan-Meier method as previously described ^30^.

### Flow cytometry analyses

C5ar1^-/-^ and WT mice were euthanised as per APLAC guidelines. The whole small intestine was washed in ice-cold PBS and the tissue was minced into 1 mm pieces. Spleen and small intestine tissue was digested into a single suspension using the murine tumour dissociation kit from Miltenyi Biotech (Auburn, CA) as per the manufacturer’s protocol. After RBC lysis, cells were re-suspended in PBS, counted and then stained with Zombie NIR (BioLegend, San Diego, CA) for live/dead cell discrimination. Nonspecific binding was blocked using an anti-mouse CD16/32 (BioLegend, San Diego, CA) antibody. Following which, cell surface staining was performed using fluorophore conjugated anti-mouse CD45.1 (30-F11), CD11b (M1/70), CD11c (N418), Ly6G (1A8), Ly6C (HK1.4), F4/80 (BM8), CX3CR1 (SA011F11), CD8 (53-6.7), CD4 (GK1.5) from Biolegend (San Diego, CA). After surface staining, cells were washed, fixed and permeabilised using BD Cytofix/Cytoperm kit and then stained for IL-10 expression (JES3-19F1, Biolegend). Flow cytometry was performed on LSR Fortessa (BD Biosciences) in the Radiation Oncology Dept. FACS facility and analysed with FlowJo software (Tree Star Inc., San Carlos, CA). Compensations were attained using Anti-Rat and anti-hamster compensation beads (BD Biosciences). For fixable live/dead staining, compensation was performed using ArC amine reactive compensation beads (BD Biosciences).

Gating schemes of dissociated tissues was as follows: Epithelial cells (ZNIR−CD45−), Immune cells-(ZNIR−CD45+), CD8 T-cells (ZNIR−CD45+CD8+), macrophages (ZNIR−CD45+CD11b+F4/80+). For positive staining determination for IL-10 and CX3CR1, respective FMO controls were used.

### Spatial transcriptomics

The Nanostring GeoMx Digital Spatial Profiler (DSP) workflow was performed on FFPE slides of mouse small intestine samples representing the following conditions, 1) WT + IgG antibody, 2) WT + IL-10 blocking antibody, 3) C5ar1^−/−^ + IgG antibody and 4) C5ar1^−/−^ + IL-10 blocking antibody. Small intestine samples from three individuals were used for each of the four conditions, selecting 12 regions of interests (ROIs) per condition, with a total of 48 ROIs selected for the DSP workflow. ROI segmentation classed spatially distinct small intestinal crypts and LGR5+ regions. Raw FASTQ and read counts data obtained have been deposited in EBI ArrayExpress, accession number: E-MTAB-15863.

Downstream analysis of the raw read counts was performed in R (ver. 4.3.2). ROI quality control (QC) and counts filtering were performed as described in the *standR* package (ver. 0.99.12) documentation. ROIs with nuclei count of less than 20 and genes expressing raw count values of <=5 in 90% of the ROIs were removed for downstream analysis^31^. Normalisation and downstream differential gene expression analysis was performed with the *DESeq2* package (ver. 1.38.3)^33^. Genes ranked by Wald statistic from the differential gene expression results were then used to perform pairwise gene set enrichment analysis (GSEA) using the *clusterProfiler* package (ver. 4.18.1) with the Gene Ontology ‘biological processes’ signature gene sets. Gene set enrichment results were then ranked by their normalised enrichment scores for assessment of the top enriched pathways in each comparison^35^.

### Cells expressing C5aR1-regulated GOIs

GSEA of Spatial transcriptomics identified significant C5aR1-regulated genes in mouse SIs. The genes were studied on the Mouse Cell Atlas MCA 2.0 dataset. Filters enabled **A.** adult mouse intestine-specific gene expression clusters to be mapped to cell-specific clusters to identify **B.** the cell types positive for a GOI.

### Spatial deconvolution analysis

Downstream deconvolution analysis of the spatial transcriptomics data utilised the *immunedeconv* (ver 2.1.0) R package^37^. The mMCP-counter algorithm outline in the package’s documentation or SeqImmuCC was applied to the spatial transcriptomics data to derive estimate immune cell fractions^38,39^.

### Macrophage maturation and ImmGen signatures

Signatures used for comparison between the different conditions were gene as described in^40^ and the Immunological Genome Project (ImmGen) respectively. Box plot visualisation of signature comparisons utilised the variance stabilised transformed unfiltered version of the spatial transcriptomics gene count data. Variance stabilising transformation of the expression matrix was performed using the *vst* function described in the DESeq2 package

### Single-cell sequencing data

The scRNA-sequencing (scRNA-Seq) data from^32^ as obtained from the Gene Expression Omnibus (GSE165318) was used. It comprised single cell transcriptomes from intestinal tissues 1 day (n = 4), 3 days (n = 3), 7 days (n = 3) and 14 days (n = 3) after a single RT dose of 10 Gy, and unirradiated tissues (n = 4). Single cells were isolated by 10X Genomics and sequenced using Illumina HiSeq 3000. Data from the unirradiated and 3 days after RT timepoint were analysed. Raw gene expression matrices and metadata were processed using the Seurat package (v5.3.0) in R (v4.3.3) according to published pipelines^34^. QC measures excluded low-quality cells expressing 200 < and > 2, 500 genes. Mitochondrial gene content was < 5 % in all analysed cells. After QC, a total of 4, 531 and 6, 773 cells, and 17, 254 and 16, 585 genes remained in the unirradiated and 3 days post-RT timepoint samples, respectively. Seurat’s *NormalizeData* and *ScaleData* functions removed variation resulting from total cellular read count and mitochondrial read count via log-normalisation and scaling. For data integration, *FindVariableFeatures* identified genes with high variability in each sample, and these were used to integrate the data via the Seurat-CCA method. Principal component analysis was performed to compute the top 20 principal components, followed by Uniform Manifold Approximation and Projection (UMAP) for visualization of cell clusters in two dimensions. Louvain clustering algorithm compared shared nearest neighbours to group transcriptionally similar cells. Cell clusters were annotated based on marker gene expression from literature studies (see table below). Marker gene signatures were scored using the UCell package (available from https://github.com/carmonalab/UCell). Dot plots were generated using ggplot2 (v3.5.2)^36^

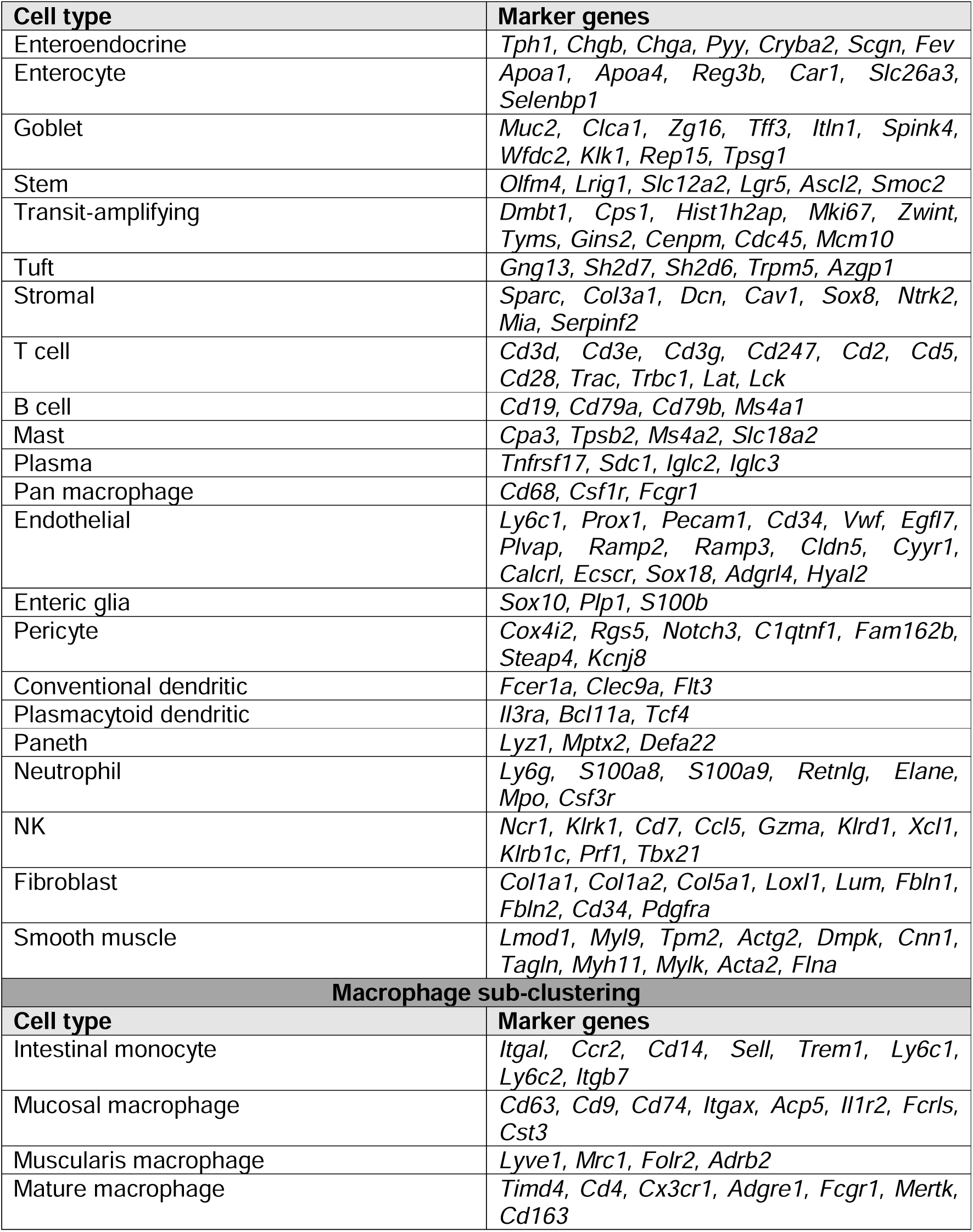

## Author contributions

Conceptualization, M.M.O. Methodology, M.M.O., H.C., I.C., S.M., D.K.M., R.K.K., Y.J., K.M.C. I.Z., and D. M., Writing original draft, M.M.O. Writing review and editing, M.M.O., H.C., and I.Z. Investigation, .M.O., H.C., I.C., S.M., D.K.M., R.K.K., and Y.J. Formal analysis, M.M.O., H.C., I.C., S.M., D.K.M., R.K.K., Y.J., and K.M.C. Resources, T.M.W. Supervision M.M.O., S.J.A.B., and A.J.G. Funding acquisition, M.M.O., A.J.G., E.G.G., and Q.T.L.

## Acknowledgements

This work was supported by the Medical Research Council (MC_UU_00001/10); NIH Grants CA-67166, CA-197713 and CA257907-01A1, the Silicon Valley Foundation, the Sydney Frank Foundation and the Kimmelman Fund (AJG). MMO was a Cancer Research Institute Irvington Fellow supported by the Cancer Research Institute. MMO is supported by the Gray Trust, and an EU Horizon Europe Research and Innovation program Cancer Mission “HIT-GLIO” (101 136 835). MMO is also supported by the Academy of Medical Sciences (AMS), the Wellcome Trust, the Government Department of Business, Energy and Industrial Strategy (BEIS), the British Heart Foundation and Diabetes UK through an AMS Springboard (REF:SBF008/1156) and Cancer Research UK (CRUK) grant number C5255/A18085, through the Cancer Research UK Oxford Centre. RKK was supported by Stanford ChEM-H Undergraduate Scholars Program.

---

**Supplementary Figure 1. Targeting C5aR1 reduces histological damage and improves survival following RT**

**(A)** Graph shows the % of LygC^+^C5aR1^+^ cells found in the spleen of WT or C5ar1^-/-^ mice. Individual points represent individual mice per group. * = p<0.05, 2-tailed t-test.

**Supplementary Figure 2. Macrophage maturation and protective radiation responses**

**(A)** Representative image of Ki67+ staining in WT irradiated (9 Gy) mice.

**(B)** Graph shows the % Ki67+ cells in WT or C5ar1^-/-^ mice irradiated with 9 Gy total abdominal irradiation. Intestines were harvested 3 days post-IR. Individual points represent individual mice per group.

**(C)** Graph shows the % of Ki67+ cells in crypts of vehicle or PMX205 treated mice irradiated with 9 Gy total abdominal irradiation. Intestines were harvested 2 days post-IR. Individual points represent individual mice per group.

**(D)** Estimated Cell fractions following mMCP-counter deconvolution of spatial transcriptomics data from Figure 2A.

**(E)** Estimated Percentage composition following SeqImmuCC deconvolution of spatial transcriptomics data from Figure 2A.

**(F)** Gating strategy example

**(G)** Graph shows the % of CD45^+^ cells found in small intestine of WT or C5ar1^-/-^ mice. Individual points represent individual mice per group.

**(H)** Graph shows the % of CD8^+^ cells found in small intestine of WT or C5aR1^-/-^ mice. Individual points represent individual mice per group. * = p<0.05, 2-tailed t-test. Individual points represent individual mice per group.

**(I)** Graph shows the percentage of CD11b^+/^F480+ macrophages in the intestine of mice treated with either IgG or anti-CSF1R antibody. ** = p<0.01, 2-tailed t-test. Individual points represent individual mice per group.

**Supplementary Figure 3. C5aR1 signalling prevents apoptotic cell clearance through an attenuated IL10-dependent response**

**(A)** Graph shows the % of CD45^-^IL10^+^ cells found in small intestine of WT or C5ar1^-/-^mice. Individual points represent individual mice per group.

**(B)** Graph shows the % of F4/80^+^IL10^+^ cells found in the spleen of WT or C5ar1^-/-^ mice. Individual points represent individual mice per group.

**(C)** Graph shows the % of CD4^+^IL10^+^ cells found in small intestine of WT or C5aR1^-/-^mice. Individual points represent individual mice per group.

## Notes

### Competing Interest Statement

The authors have declared no competing interest.

