## Supplementary figures and images for "Targeting C5aR1 Reveals Protective Macrophage Maturation States in Intestinal Injury"

### Supplementary Figures

**A**

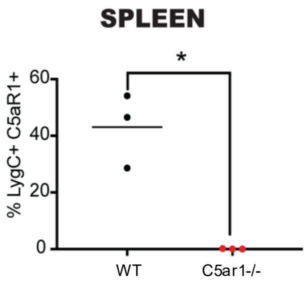

#### Supplementary Figure 2

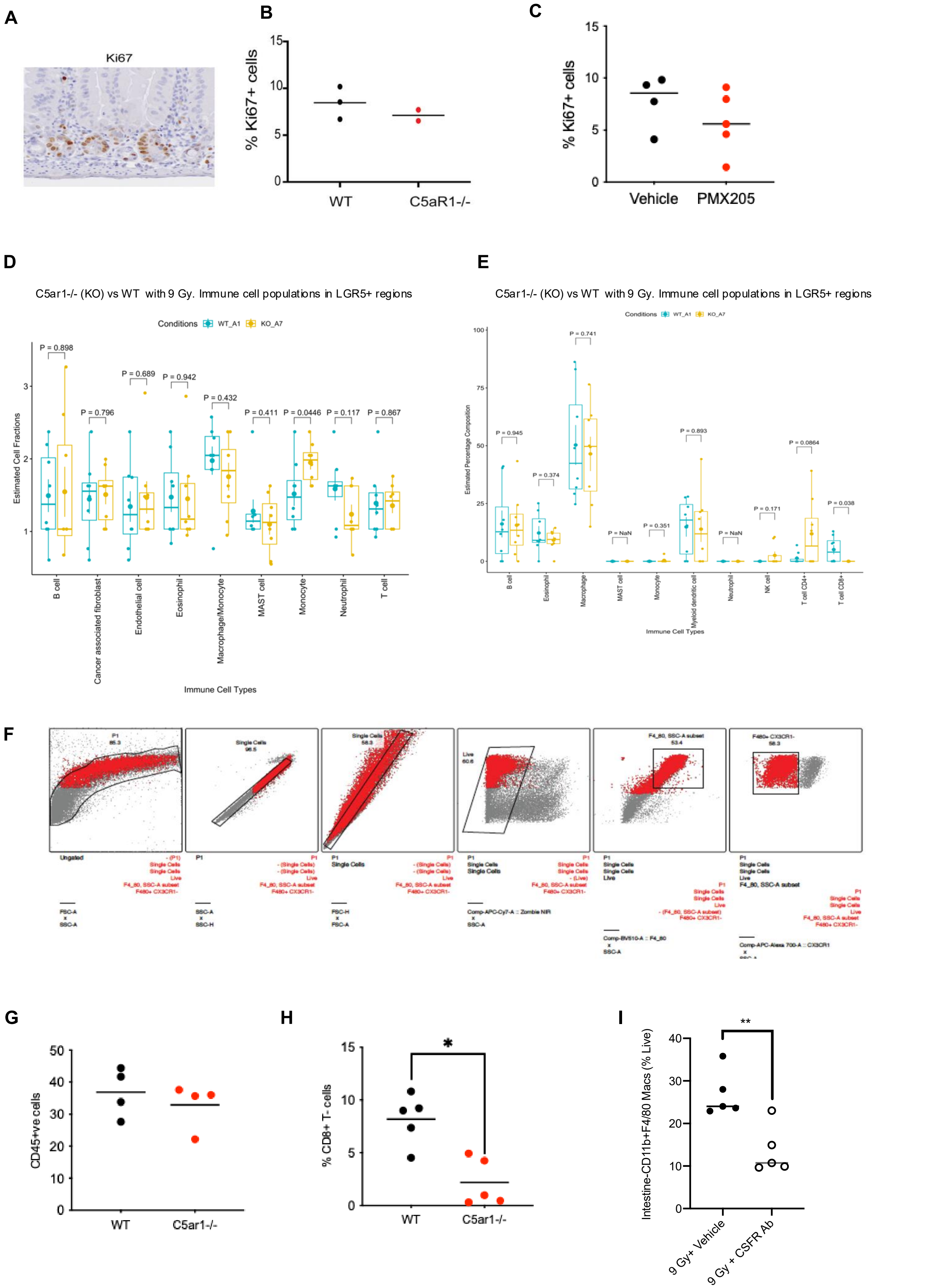

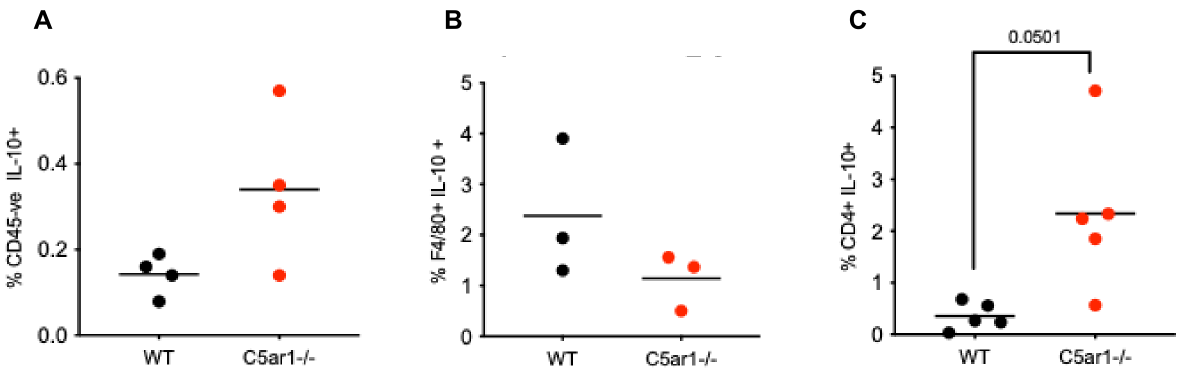
